# Graph Neural Networks Ameliorate Potential Impacts of Imprecise Large-Scale Autonomous Immunofluorescence Labeling of Immune Cells on Whole Slide Images

**DOI:** 10.1101/2022.08.28.505606

**Authors:** Ramya Reddy, Ram Reddy, Cyril Sharma, Christopher Jackson, Scott Palisoul, Rachael Barney, Fred Kolling, Lucas Salas, Brock Christensen, Gregory Tsongalis, Louis Vaickus, Joshua Levy

## Abstract

The characteristics of tumor-infiltrating lymphocytes (TIL) are essential in cancer prognostication and treatment through the ability to indicate the tumor’s capacity to evade the immune system (e.g., as evidenced by nodal involvement). Machine learning technologies have demonstrated remarkable success for localizing TILs, though these methods require extensive curation of manual annotations or restaining procedures that can degrade tissue quality, resulting in imprecise annotation. In this study, we co-registered tissue slides stained for both hematoxylin and eosin (H&E) and immunofluorescence (IF) as means to rapidly perform large-scale annotation of nuclei. We integrated the following approaches to improve the prediction of TILs: 1) minimized tissue degradation on same-section tissue restaining, 2) developed a scoring algorithm to improve the selection of patches for machine learning modeling and 3) utilized a graph neural network deep learning approach to identify relevant contextual features for lymphocyte prediction. Our graph neural network approach accounts for surrounding contextual micro/macro-architecture tissue features to facilitate interpretation of registered IF. The graph neural network compares favorably (F1-score=0.9235, AUROC=0.9462) to two alternative modeling approaches. This study brings insight to the importance of contextual information leveraged from within and around neighboring cells in a nuclei classification workflow, as well as elucidate approaches which enable the rapid generation of large-scale annotations of lymphocytes for machine learning approaches for immune phenotyping. Such approaches can help further interrogate the spatial biology of colorectal cancer tumors and tumor metastasis.

## 1. Introduction

Numerous studies have demonstrated that immune cell infiltrates play a crucial role in the adaptive immune response for specific bacterial and viral infections and various types of cancers (Aoshi et al., 2011; Galon et al., 2006). In the context of cancer, the presence, type, and location of tumor-infiltrating lymphocytes (TIL) play a crucial role in prognostication as this can be indicative of the tumor’s capacity to evade or suppress the immune system (Morrison et al., 2022; Whiteside, 2022). It is believed that TIL at the primary site may indicate concurrent or future activity at the regional lymph node (Caziuc et al., 2019). As such, localizing TILs at the primary site at the time of resection may indicate that a less invasive secondary treatment is required (e.g., adjuvant chemotherapy, radiotherapy).

Many methods for determining the presence of TIL lack spatial resolution. However, hematoxylin and eosin (H&E) or immunohistochemical (IHC) stained tissue slides allow for spatial assessment. H&E provides a morphological and cytological examination, and IHC allows for multiplexing of protein markers that can disaggregate distinct cellular populations. For instance, immunoscore is a digital pathology technology that can assess the density of CD8+ (cytotoxic) and CD3+ (co-receptor which activates cytotoxic T cells) T cells inside the tumor and at the invasive margin and is highly effective for prognostication (Kwak et al., 2016). Increased density is associated with lower progression-free survival as the presence of these cells can signal immune exhaustion, though alternatively could be associated with favorable prognosis through induced anti-tumoral cytotoxicity (Bruni et al., 2020). Studying similar effects using routine H&E stained slides through accurate localization of these immune cells is an emerging study area. Inferring locations of TIL is particularly challenging because it either requires large-scale annotation or wash and restain procedures to tag H&E stained tissue with various protein markers which deforms and degrades tissue. However, several emerging applications for careful restaining have demonstrated success in tagging millions of cells with molecular information with relatively little effort (Jackson et al., 2020b). Tagging with IHC can also be done using serial sections, but is suboptimal as there is no microarchitectural alignment due to the 5-micron separation between adjacent levels. Immunofluorescence (IF) staining can label several antigens in the same slide through the emission of relatively discrete imaging spectra (which has higher multiplexing potential compared to IHC). Furthermore, for tagging multiple markers in addition to routine staining, the application of IF after H&E is arguably less destructive than destaining H&E then staining with IHC on the same section. Several methods have been proposed to infer IF computationally, though, these generally rely on serial section staining (Burlingame et al., 2018). For the application of H&E after IF, registration can still complicate analyses with imprecise alignment though it is preferred to serial staining as it maintains significantly better microarchitectural alignment.

Machine learning algorithms, in particular, deep learning through the use of artificial neural networks (ANN), have demonstrated remarkable performance across a wide variety of image classification and detection tasks, all of which are relevant for atomizing lymphocytes for further analysis. Convolutional neural networks (CNN), a deep learning approach that accounts for spatial dependencies in images have gained increased attention over the past few years for TIL-specific inference tasks. Notable methods include segmentation (e.g., U-Net) (Jackson et al., 2020a; Saltz et al., 2018; Turkki et al., 2016), detection (i.e., panoptic segmentation, Fast R-CNN, Panoptic FPN; as popularized by the Detectron2 library (Wu et al., 2019)), and generative adversarial network approaches (Burlingame et al., 2018). Relatively unexplored is the use of detection networks (e.g., Detectron2) for these prediction tasks. Furthermore, surrounding spatial information can provide additional context.

In this study, we aim to explore algorithmic methods which, when used in conjunction with IF staining, can predict the presence of TIL while remaining sensitive to the imprecision in H&E cell-tagging from microarchitectural registration. We hypothesize that nascent graph neural network deep learning methods for cell type inference based on neighboring cells and micro/macro-architecture, can ameliorate lymphocyte inference challenges associated with IF tagging. Furthermore, we attempt to improve selected datasets for prototyping our algorithm through a patch-wise registration scoring algorithm.

Here, we investigate the effectiveness of graph neural networks (GNN) in identifying lymphocytes from H&E that were tagged through imprecisely registered IF.

## 2. Methods

### 2.1. Methods Overview

We performed: 1) large-scale annotation of lymphocytes using IF stains from the same section as the H&E, 2) developed a scoring algorithm to improve the selection of patches for algorithmic prototyping, and 3) utilized a GNN to mine contextual features relevant for IF-guided lymphocyte prediction. In brief, our method is as follows (See **Appendix A**):

1. Acquire H&E and IF stained whole slide images (WSI) from the same tissue section from 36 stage III matched colorectal cancer patients at Dartmouth Hitchcock Medical Center **(Data Collection, Section 2.2, Figure 1A)**.
2. Train U-net and detection neural network models on pathologist annotations from a combination of external public and private datasets to infer pixel-wise presence of nuclei and localize nuclei instances respectively **(Nuclei Detection, Section 2.3, Figure 1B)**.
3. Perform patch-wise registration of IF patches to H&E sections by aligning SYTO and Hematoxylin stains **(Stain Registration, Section 2.4)**.
4. Simultaneously score registration quality for specific patches and identify the optimal image intensity of the CD45 stain, to tag immune cells, on the H&E using a sensitivity analysis comparing the SYTO stain to the U-Net results **(Alignment Screening, Section 2.4, Figure 1C)**.
5. Use the Detectron2 nuclei detection model to create an annotated cell dataset from the stained WSIs, containing information on where the cell was located and whether it was an immune cell **(Cell Tagging, Section 2.5, Figure 1D)**.
6. Using the tagged cells, train and compare Detectron2, CNN, and GNN models for their ability to detect lymphocytes based on the annotated cell dataset **(Model Training, Section 2.6, Figure 1E)**.

**Figure 1:**
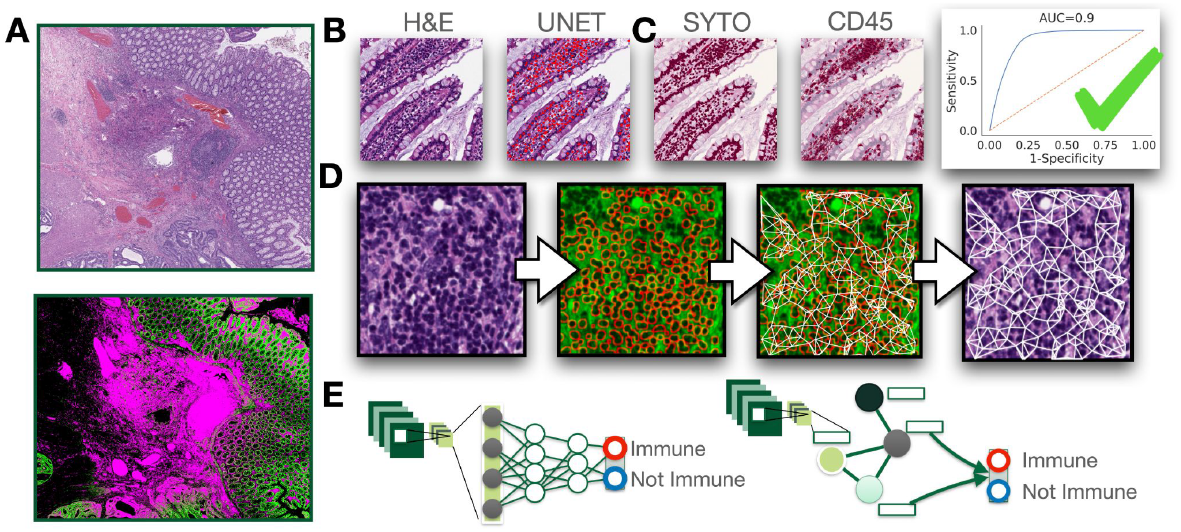
Workflow for cell dataset generation for lymphocyte prediction models: A) H&E and IF stains are collected and coregistered; B) UNET trained to predict nuclei to screen out C) slides based on concordance with SYTO stain via sensitivity analysis; D) graphical representation of detectron prediction and cell graph generation; E) application of CNN and GNN modeling approaches for immune cell prediction

### 2.2. Data Collection

The primary dataset utilized in this study was acquired from 36 Stage-III matched (pTNM system) colorectal cancer patients at Dartmouth Hitchcock Medical Center, determined through a retrospective review of pathology reports from 2016 to 2019 following IRB review and approval. Half of the patients had concurrent tumor metastasis and were otherwise matched on age, sex, tumor grade, tissue size, mismatch repair status, and tumor site using iterative patient resampling with t-tests for continuous variables and fisher’s exact tests for categorical variables. Tissue blocks were sectioned into 5-micron thick layers. Sections were stained with fluorescent-labeled, IF, antibodies for the following markers: 1) tumor/epithelial (PanCK), immune cells (CD45), and nuclei (SYTO13). These IF stains were initially acquired for a previously published study on spatial immune markers of metastasis, which utilized the GeoMX Digital Spatial Profiler (DSP, Nanostring Technologies, Seattle, WA) for image scanning into 16-bit unsigned color (one channel per stain) TIFF format images (Levy et al., 2022). After IF staining, the same sections were stained for H&E (without requiring destaining as the chemical reagents of the H&E minimally interacted with the fluorophores) and scanned into WSI using the Aperio AT2 scanner at 20x (8-bit unsigned color). The DHMC in-house dataset consisted of 36 WSIs, divided into 6,654 subarrays. Each of the subarrays were 768 pixels in each spatial dimension for patch-wise alignment and were further divided into nine square subarrays of side length 256-pixels without overlap, resulting in a total of 59,886 subarray images for cell identification.

Separately, we assembled an in-house dataset of 2,155 pathologist-annotated nuclei and a publicly available dataset of 30,837 pathologist-annotated nuclei to develop initial nuclei segmentation and detection approaches (Kumar et al., 2017, 2020).

### 2.3. Initial Nuclei Segmentation and Detection Models for Cell Localization

First, using the assembled nuclei detection dataset, a U-Net model was trained to detect the hematoxylin-stained nuclei on a pixel-wise basis for alignment scoring. The Detectron2 nuclei detection model was also trained on the same dataset which was more sensitive to adjacent cell boundaries through the adoption of panoptic segmentation methods and better allowed for cell counting (Wu et al., 2019).

The nuclei detection model was pre-trained with a 3x schedule that was available through the public Detectron2 Model Zoo, and it was then trained on the in-house data for a maximum of 5,000 epochs. Training was stopped when overfitting occurred and when accuracy on the test set crossed 90%. The base learning rate was set to 0.0125 and 5 images were used per iteration. The model used a Mask R-CNN architecture that has a Residual Network+Feature Pyramid Network (ResNet+FPN) backbone based on the ResNet-101 model. It was tested using a detection threshold of 0.05 and a non-maximum suppression (NMS) threshold of 0.25. All hyperparameters were set to the Detectron2 config defaults if they were not otherwise specified.

### 2.4. Slide registration and screening imprecise alignments through sensitivity analysis

H&E and IF WSIs were registered through patch-wise alignment algorithms applied to the nuclear stains, Hematoxylin (determined using the Macenko stain deconvolution method), and SYTO13 staining intensities respectively. As the tissue was minimally deformed during H&E staining, the H&E and IF sections from the same tissue specimen were co-registered through low-resolution rigid transformations. Then, both the H&E and IF WSI were divided into 768-pixel patches for more precise microarchitectural alignment (**Appendix A.1**). After registering the IF and H&E nuclear stains, the CD45 stain was overlaid by leveraging the same displacement field as the SYTO13 stain to tag immune cells.

We employed several mechanisms to screen out poorly aligned tissue patches. First, we calculated the pixel-wise difference in normalized staining intensities between the nuclear stains. The patch was removed from the set if this difference exceeded a specific threshold (mean squared error of 80). We also applied the trained U-Net model on the H&E patches to establish nuclei annotations. For each patch, we used a sensitivity analysis to calculate a C-statistic to provide an overall measure of agreement between the IF nuclear stain intensity and the predicted nuclei mask across a range of intensity thresholds. Patches with an overall agreement C-statistic of at least 0.85 were included in the set **(Figure S1)**. The sensitivity analysis was also used to identify a staining intensity threshold used to establish an IF nuclear mask based on maximum fidelity to the nuclei mask predictions. The same intensity threshold was applied to the CD45 stain to establish an immune cell mask, which was confirmed through visual inspection with collaborating pathologists.

### 2.5. Lymphocyte Prediction Dataset

There were no initial pre-existing annotations of immune and non-immune cells for the lymphocyte prediction model. By leveraging a highly accurate nuclei detection model (**Appendix A.2**) and registered immune cell masks, we were able to detect and label 5,377,681 nuclei, after filtering false positives. Nuclei were algorithmically annotated by overlaying the immune cell masks (**Appendix A.1**). This dataset contained 953,274 immune cells and 4,424,407 non-immune cells in the training/validation set and 19,408 immune cells and 90,231 non-immune cells in the test set.

### 2.6. Lymphocyte Prediction Modeling Approaches

We compared the following model approaches for the prediction of immune cells across our newly annotated dataset: 1) a cell detection model (Detectron2 framework) which outputs two classes (immune/non-immune), 2) a CNN-trained off of small images extracted around the bounding boxes, and 3) a GNN model trained on embeddings extracted from the CNN. Details of each model specification can be found below (**Appendix A.3, A.4**).

Detectron2, unlike other methods, can automatically detect the lymphocyte cells from the original image instead of just classifying subimages like the CNN and GNN.

We trained a convolutional neural network (i.e., ResNet18) on 64-pixel patches extracted around the initially predicted nuclei as means to more precisely model cellular morphology. Embeddings of the cells were extracted from the penultimate layer of the CNN, as well as the cells’ positional x-y coordinates. These coordinates were used to generate graph datasets using both a k-nearest neighbor and radius neighbors graph for the GNN to train on. Cells represent nodes of the graph and were connected by local proximity. Node attributes were captured using CNN embeddings.

A GNN model was used to integrate macroarchitectural cues for the inference of immune and non-immune cells (Ahmedt-Aristizabal et al., 2022; Fey and Lenssen, 2019). Such a model could potentially help overcome imprecise labeling from the automated registration, filtering, and application of the co-registered immune cell mask. The model outputs a vector representing the likelihood of observing an immune cell **(Figure S2)** (**Appendix B**,**C**).

### 2.7. Model evaluations

The classification performances of the lymphocyte prediction models were determined through reports of the accuracy, F1-score, area under receiver operator curve (AUROC), and intersection over union (IOU). A F1-score is equal to the harmonic mean of the precision and recall, and was calculated through the Scikit-learn Python library. Due to the inherent class imbalance between immune and non-immune cells in the data, we considered the weighted F1-score, with thresholds calculated using Youden’s index (Ruopp et al., 2008). We depicted the number of true positives, true negatives, false positives, and false negatives through a confusion matrix, where immune cells were considered positive classifications and non-immune cells were considered negative classifications for calculation of sensitivity and specificity statistics. All of the previously described methods were used to compare the Detectron2, CNN, and GNN models. IOU was only used for the Detectron2 model, as it was the only model that had an output of bounding boxes, and showed how the predicted classifications compared to the annotated classifications. The CNN and GNN models were also interpreted through the generation of Uniform Manifold Approximation and Projection (UMAP) (McInnes et al., 2018) embeddings of the nuclei, which allowed for visual assessment on the ability of the neural networks to delineate cell types.

## 3. Results

All three modeling approaches achieved an F1-score above 0.8 (**Appendix D**). Both the CNN and the GNN models outperformed the cell detection model for their ability to delineate immune cells. Notably, the GNN obtained optimal performance after taking into account Youden’s threshold, obtaining an F1-score of 0.92 **(Table 1, Figure S3)**.

**Table 1:** Summary of model classification metrics

| Model | Accuracy | F1-Score | AUROC | IOU |
| --- | --- | --- | --- | --- |
| Detectron2 | 82.11 | 0.8175 | 0.6961 | 0.6737 |
| CNN | 90.35 | 0.8995 | 0.9297 | N/A |
| GNN | 92.45 | 0.9235 | 0.9462 | N/A |

We visualized the UMAP projection of the extracted CNN and GNN embeddings of cells from the WSI **(Figure S4)**. The GNN model was able to learn contextual features and delineate cell types based on co-localized cells that gave this model a competitive advantage over the CNN model. We also visually compared the output from all three modeling approaches versus the ground truth on a few randomly selected patches which corroborated with the aforementioned findings **(Figure 2)**.

**Figure 2:**
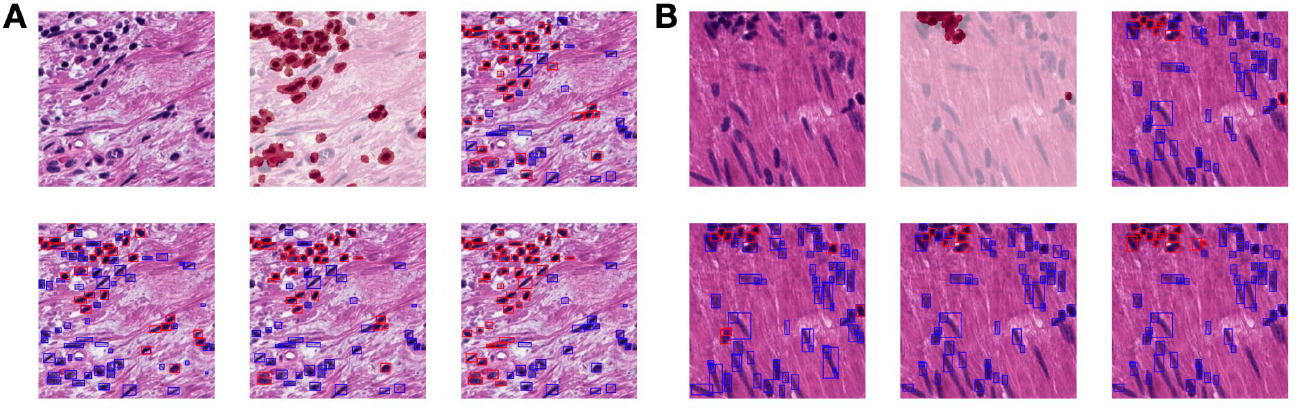
WSI patch, immune mask, ground truth images on upper row with predictions from Dectron2, CNN, and GNN models on bottom row for each example image subarray (A-B)

## 4. Discussion

The tumor immune microenvironment is an amalgamation of immune cells, chemokines, cytokines, and other immune modulators and plays a crucial role in coordinating the immune response to processes governing tumorigenesis and metastasis. As such, understanding spatial biology at the primary site is crucial for informing timely and relevant disease management options. Thus, the localization and quantification of distinct immune cell lineages may help inform the development of new spatial biomarkers. Informatics methods are still being developed to make sense of the data from this nascent field. Inference from morphological findings from an H&E tissue slide is an attractive approach because H&E staining is routinely done and inexpensive. However, optimal means of data collection and annotation are presently quite onerous and are an active area of exploration. As pathologists may incorrectly localize immune cells, IF staining to tag cells may present a viable alternative for labeling at scale, though it is expected that detection networks may struggle to make use of the antigenically tagged information.

In this study, we detailed an approach for the rapid and accurate immune cell annotation of nuclei based on the registration of IF which requires no tissue destaining. We applied geometric deep learning methods to potentially ameliorate inexact cell tagging by explicitly leveraging morphological and architectural information from neighboring cells. Our preliminary analysis suggests that GNNs, when combined with IF tagging of nuclei can accurately localize and tag immune cells in WSIs. We plan to further investigate the clinical utility of this technique and will further iterate and improve this method for downstream approaches for large-scale phenotyping in the context of studying tumor metastasis.

There were some limitations in the methodology of this study. We assumed the initial nuclei detection model achieved sufficient accuracy as judged by visual inspections from our practicing pathologists. While there may have been some inaccuracies in the initial cell localization, this is not outside of what is expected from other similar studies which leverage these datasets and further exploration is outside of the study scope (Mahmood et al., 2020). Furthermore, the accuracy of our approach may have been impacted by manual staining processes for IF and batch effects. For instance, some slides were coverslipped for a few days, which may have impacted the specificity of the IF stain but was a pragmatic consideration when planning our experiment due to technical staffing shortages. Staining for H&E, by contrast, used automated staining protocols. Staining and cell tagging inaccuracies may have also been introduced in the registration process though we attempted to control for this through the sensitivity analysis. Future iterations of this model will attempt to more tightly control experimental preplanning through additional workflow automation.

Despite these limitations, our proposed lymphocyte prediction tool is valuable for researchers aiming to study spatial biology as it allows for the easy creation of robust IF tagged dataset on millions of cells even in small-scale clinical feasibility studies. When pairing with macroarchitectural annotations (e.g., within the tumor, at the tumor-immune interface, etc.) identifying immune cells in these regions can help infer the impact of immune cells in these regions for outcome such as metastasis, recurrence and survival. Furthermore, the presence of different cell types can confound or reduce the power of molecular association studies on microdissected tumor (Aran et al., 2015). Several recent studies have explored machine learning-based inference of highly multiplexed protein and RNA markers inferred on the cellular/subcellular level (He et al., 2020; Moses and Pachter, 2022; Zeng et al., 2022). Integrating histological information with predicted cell types through deconvolution approaches can more precisely identify canonical cellular populations (e.g., FOXP3+ T regulatory cells) which could further inform the coordinated response. In the future, we aim to investigate the interplay between histomorphology and protein/RNA expression localized to distinct locations on the slide through the adoption of highly multiplexed spatial assays including the GeoMX DSP and 10x Genomics Visium Spatial Transcriptomics.

## 5. Conclusion

In this work, we demonstrated the first application of GNN methods to H&E slides that were tagged through co-registered IF in the context of studying TILs. Our study suggests that contextual information leveraged from neighboring cells are important for nuclei classification and this workflow, as a whole, can be effective for generating large-scale immune phenotype data for studying the spatial biology of colorectal cancer tumors and tumor metastasis. We plan to further standardize this process and employ measures to explore the downstream implications of these findings with high precision.

## Acknowledgments

This work was supported by the Dartmouth Hitchcock Medical Center Department of Pathology and Laboratory Sciences through the Emerging Diagnostic and Investigative Technologies program.

## Appendix A. Supplementary Methods

### A.1. Microarchitectural Alignment and Additional Cell Tagging Details

For each pair of nuclear stains, akaze features were extracted from each nucleus stain image from matching patches, and a k-nearest neighbors and radius neighbors brute force feature matcher were used to identify matching local features between the images. Matched features between the H&E and IF nuclear stains were used to compute a perspective transformation for the final registration.

The immune cells were tagged by calculating the percentage overlap between the predicted nuclei instance mask/bounding box by the cell detection model and the immune cell mask. If at least 25% of the nucleus instance mask was labeled pixel-wise as an immune cell, the nucleus was tagged as such and was not labeled as an immune cell if it failed to surpass this threshold.

**Figure S1:**
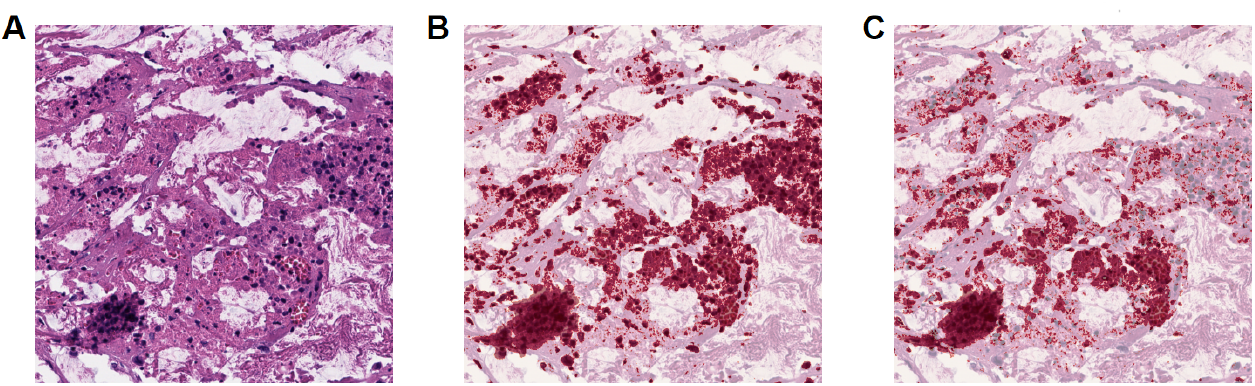
Comparison of (A) H&E stain on a WSI with (B) an additional nuclei mask (SYTO13) and (C) immune mask (CD45)

### A.2. Detection Dataset Format

These datasets were prepared in the Microsoft COCO (MS COCO) format. The COCO format is commonly used for machine learning and computer vision projects and can be used for object detection, segmentation, and captioning. It uses the JavaScript object notation (JSON) format and includes information about the categories the images were classified into, raw image information such as filename and image size, and a list of all object annotations for every image in the dataset. Training and validation datasets were generated which contained nuclei annotations. For annotations, nuclei areas were calculated using the contour area method of the OpenCV-Python package based on manually segmented splines placed by the pathologists. The bounding box format for the annotations also followed the default boxmode of absolute minimum-X and minimum-Y coordinates, width, and height.

The Detectron2-derived lymphocyte prediction model dataset followed the same MS COCO dataset format as what was done for the nuclei detection model (e.g., similar area and boxmode annotations). However, there were two categories of images: immune and non-immune cells. The datasets used for the CNN and GNN lymphocyte prediction models were also adapted from the Detectron2 model datasets.

### A.3. Addressing Class Imbalances

There was also a significant class imbalance of lymphocyte and non-lymphocyte nuclei, with a ratio of approximately one immune cell for every four non-immune cells. To address these challenges, we employed class balancing techniques, such as resampling (dynamic batch-wise undersampling), reweighting the model objective, and using evaluation metrics that were relatively robust to these differences. We found these methods did not significantly impact performance. It is expected that without reweighting, the decision threshold would be shifted to reflect this proportion of immune cells.

### A.4. Additional Motivation/Training Details for Modeling Approaches

We used the Detectron2 library instead of other object detection methods like YOLO because many of the methods featured in the Detectron2 framework are easier to implement, more accurate than other methods, and had demonstrated to us favorable performance in underrepresented classes. Lymphocyte prediction relied heavily on the modeling results from our initial nuclei detection model, which generated training datasets for all three modeling approaches.

The Detectron2 lymphocyte prediction model also takes less time to train compared to the CNN and GNN models because it contains state-of-the-art prewritten libraries based on PyTorch; however, the study results indicate that the Detectron2 model requires a relatively extensive amount of data **(Table 1)**. Due to this, Detectron2 is able to make predictions more quickly during inference.

The Detectron2 lymphocyte prediction model used the same pre-trained model, the number of training images per step, and Mask R-CNN architecture as that of the nuclei detection model. The model was trained for 4,000 epochs at a base learning rate of 0.0125. Training was stopped after no significant change in validation set accuracy. Visual assessments of the model predictions were done at a detection threshold of 0.2 and NMS threshold of 0.3. All hyperparameters were set to the Detectron2 configuration defaults if not otherwise specified. The model output bounding boxes, category names, and measure of model certainty (%) of the category for each detected cell.

As a classification model, the CNN had an additional step, requiring the Detectron2 detected cell annotations to be used as an input, instead of just an image, in order to have an output of category classifications. The model capacity of the residual neural network is greater than cell class prediction layers after proposed regions of interest from Detectron2. The CNN model was trained using the PyTorch framework. A data loader was configured which loads detected cells into a Torch tensor format. Data augmentation was performed including horizontal and vertical flips, random rotations, and color jitter. The CNN lymphocyte prediction model was trained on the training dataset of nuclei for 20 epochs, a learning rate of 0.0125, and a batch size of 128 cells. The validation set F1-score was assessed after each epoch; ultimately, we saved the model at the epoch with the highest F1-score.

The GNN model had two extra steps; the first step is the same as that of the CNN model, where it requires the Detectron2 nuclei detection model to process an image, and a second step of extracting embeddings (or features) from the detected cells. The GNN model creates a more abstract representation of the WSI through the graph datasets and extracted embeddings, which can lead to an overall better generalization of the data. The model was trained for 300 epochs with a batch size of 1 and a learning rate of 1e-4 and similar to the CNN model, we kept track of the epoch which achieved the highest F1-score and saved the model when this occurred. The model was trained using the PyTorch-Geometric software framework, which takes as input a graph dataset and outputs a probability vector (Torch tensor).

**Figure S2:**
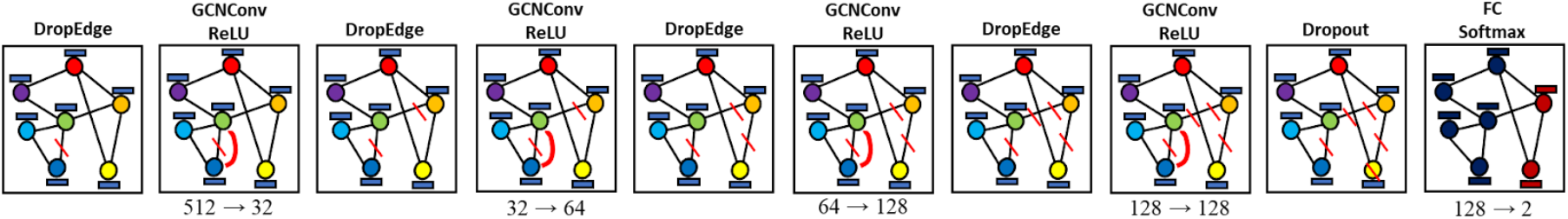
Graph Neural Network architecture

## Appendix B. Code Availability

The python programming language (Version 3.8.8) was used in all coding aspects of this study. Code was prototyped using Jupyter notebook (version 6.4.11) and leveraged computing resources (Tesla v100s GPUs) housed at the Dartmouth College Discovery Research Computing cluster. Code is available upon reasonable request.

## Appendix C. Supplementary Discussion

It should be noted that there remains outstanding debate on optimal feature extraction methods for CNN and GNNs, specifically for inferring immune cell types, which are outside of the study scope. While we employed feature extraction methods across the cells, this component has not been well explored and could shed light on what morphological and architectural information is relevant for cell typing (e.g., nuclear morphology, cytoplasm, surrounding architecture). As an example, applying GNN to large patches extracted around nuclei as opposed to nuclear morphology would support the hypothesis that context matters. In the future, we would want to experiment with more complex GNN and CNN feature extraction models for the GNN, rather than simply use a pretrained CNN model, as it could yield more nuanced information for cell type classification.

## Appendix D. Supplementary Result Figures

**Figure S3:**
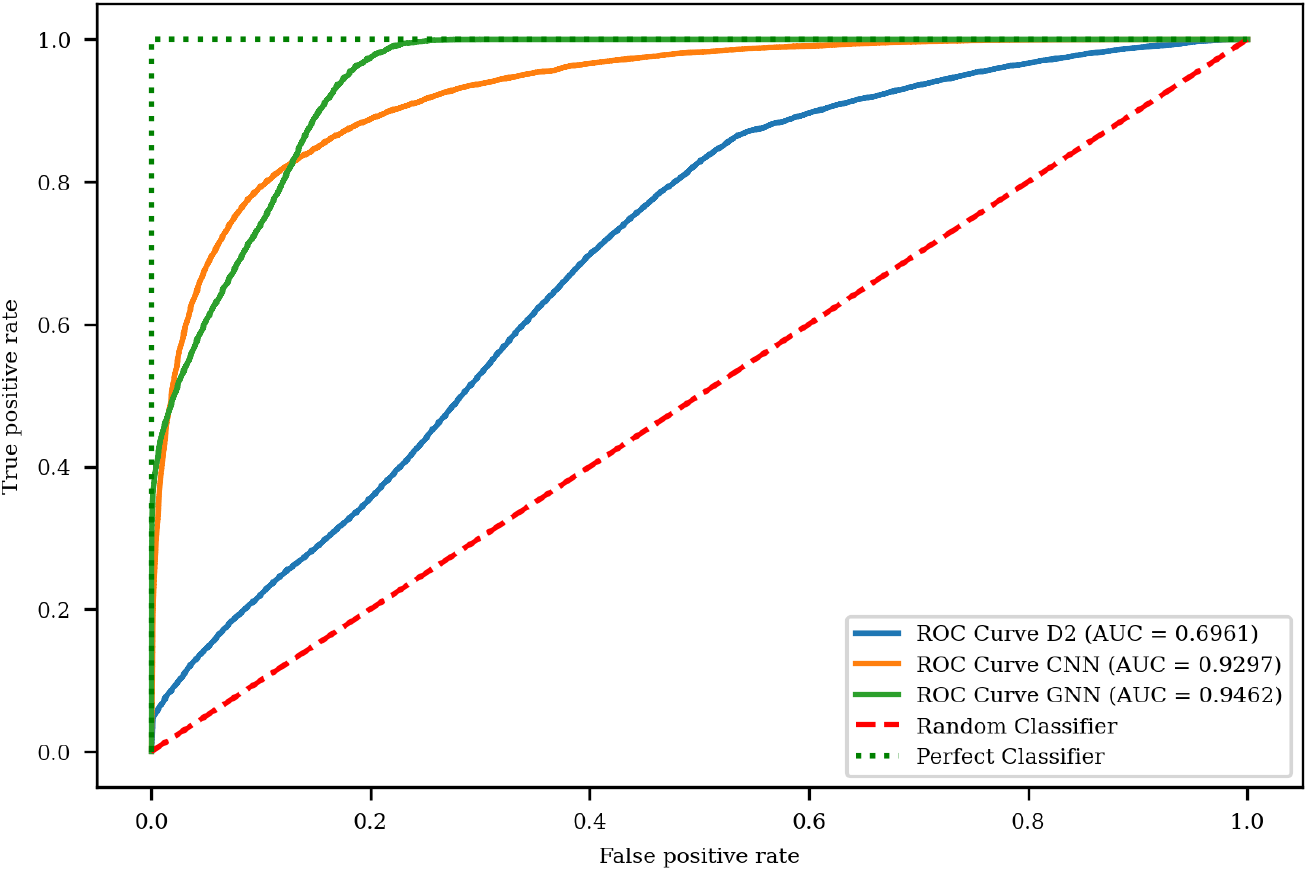
Graph comparing ROC curves of models

**Figure S4:**
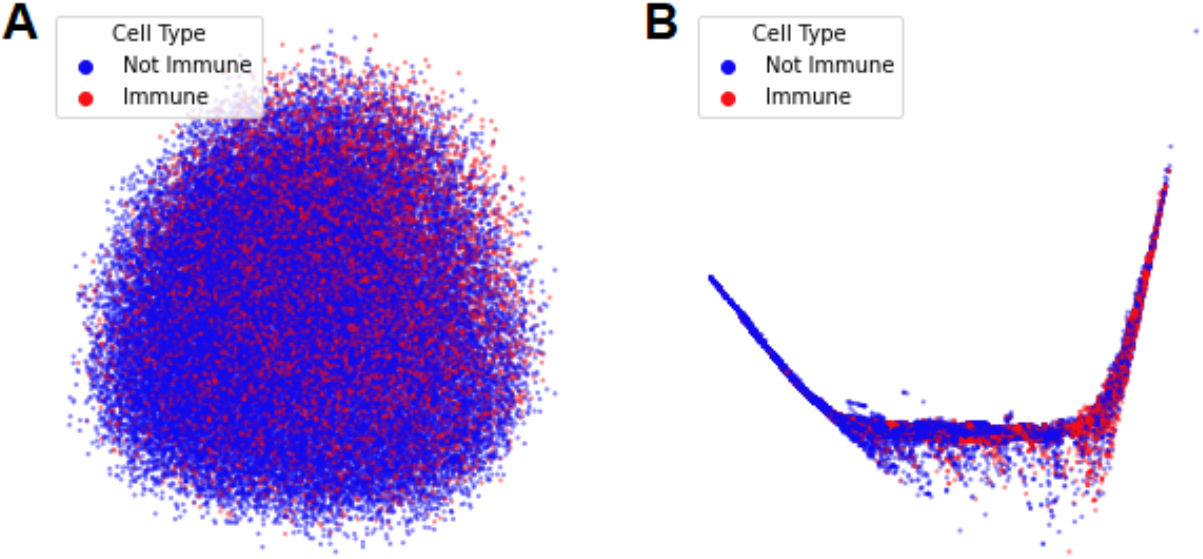
PCA projection of extracted embeddings from cell images using (A) CNN and (B) GNN models

## Notes

### Competing Interest Statement

The authors have declared no competing interest.

### Summary of Updates

Authors list updated

